# SNP-CRISPR: a web tool for SNP-specific genome editing

**DOI:** 10.1101/847277

**Authors:** Chiao-Lin Chen, Jonathan Rodiger, Verena Chung, Raghuvir Viswanatha, Stephanie E. Mohr, Yanhui Hu, Norbert Perrimon

## Abstract

CRISPR-Cas9 is a powerful genome editing technology in which a single guide RNA (sgRNA) confers target site specificity to achieve Cas9-mediated genome editing. Numerous sgRNA design tools have been developed based on reference genomes for humans and model organisms. However, existing resources are not optimal as genetic mutations or single nucleotide polymorphisms (SNPs) within the targeting region affect the efficiency of CRISPR-based approaches by interfering with guide-target complementarity. To facilitate identification of sgRNAs (1) in non-reference genomes, (2) across varying genetic backgrounds, or (3) for specific targeting of SNP-containing alleles, for example, disease relevant mutations, we developed a web tool, SNP-CRISPR (https://www.flyrnai.org/tools/snp_crispr/). SNP-CRISPR can be used to design sgRNAs based on public variant data sets or user-identified variants. In addition, the tool computes efficiency and specificity scores for sgRNA designs targeting both the variant and the reference. Moreover, SNP-CRISPR provides the option to upload multiple SNPs and target single or multiple nearby base changes simultaneously with a single sgRNA design. Given these capabilities, SNP-CRISPR has a wide range of potential research applications in model systems and for design of sgRNAs for disease-associated variant correction.

## INTRODUCTION

The CRISPR-Cas9 system, a repurposed bacterial adaptive immune system, is a powerful programmable genome editing tool for research, including in eukaryotic systems, that also has potential for gene therapy (PICKAR-OLIVER and GERSBACH 2019). With this system, *Streptococcus pyogenes* Cas9 nuclease is directed to a target site or sites in the genome that have a unique 20 nt sequence followed by a 3 bp sequence conforming to *NGG* known as the protospacer adjacent motif (PAM). A double-strand break (DSB) induced by Cas9 nuclease recruits the cellular machinery, which can repair the break either through the error-prone non-homologous end-joining (NHEJ) pathway or through homology directed repair (HDR). NHEJ often results in insertions and/or deletions (indels), which can result in frameshift mutations. HDR allows researchers to introduce or ‘knock in’ specific DNA sequences, such as precise nucleotide changes or reporter cassettes.

In addition, catalytically dead forms of Cas9 have been fused with different effector proteins to manipulate DNA or gene expression (PICKAR-OLIVER and GERSBACH 2019). For example, to correct disease-causative point mutations, CRISPR-Cas9 mediated DNA base editing has been developed as a promising method to convert undesired spontaneous point mutations to the wild-type nucleotide (GAUDELLI *et al.* 2017; KOMOR *et al.* 2016; PICKAR-OLIVER and GERSBACH 2019). DNA base editing can be achieved by fusing a Cas9 nickase with a cytidine deaminase enzyme and uracil glycosylate inhibitor to achieve a C->T (or G->A) substitution. Similarly, a transfer RNA adenosine deaminase is fused to a catalytically dead Cas9 to generate A->G (or T->C) conversion. Notably, unlike for knock-in, DNA editing-induced changes occur without a DSB and without the need for introduction of a donor template. Disease-relevant mutations in mammalian cells can be corrected with base editing strategies (DANDAGE *et al.* 2019). Prime Editing based on the fusion of Cas9 and reverse transcriptase, is another recently published technique that could add more precision and flexibility to CRISPR editing (ANZALONE *et al.* 2019). Thus, programmable editing of a target base in genomic DNA provides a potential therapy for genetic diseases that arise from point mutations.

Single-nucleotide polymorphisms (SNPs) can be defined as single-nucleotide differences from reference genomes. The targeting efficiency of Cas9 has been examined using data from genome-wide studies combined with machine learning (CHUAI *et al.* 2018; DOENCH *et al.* 2014; LISTGARTEN *et al.* 2018; NAJM *et al.* 2018; TYCKO *et al.* 2019). The position of specific nucleotides in the target sequences has been shown to affect targeting efficiency, which is the major determinant of CRISPR-Cas9 dependent genetic modification (DOENCH *et al.* 2014; HOUSDEN *et al.* 2015). Therefore, the presence of a SNP (or of an indel) can cause inefficient binding of the Cas9-sgRNA ribonucleoprotein (RNP) complex, resulting in inefficient genome editing.

Many rules for sgRNA design are generalizable and many web tools have been developed to predict sgRNA sequences for the human genome and genomes of numerous model organisms. There are two types of input that sgRNA design tools typically accept: (1) gene symbols or genome coordinates and (2) sequences. Resources that support the former typically precompute sgRNAs based on annotated reference genome information. Moreover, these sgRNA sequences are designed based on a single wild-type reference sequence without considering variants (e.g. CHOPCHOP, GuideScan; Table 1). With the second type of input, some tools BLAST user input sequence against a reference genome and correct any differences introduced by the user, thus making it impossible to design sgRNAs against a variant allele (this is the case for example for CRISPR-ERA and CRISPR-DT; Table 1). With others, it is possible to design a sgRNA to target variant allele (e.g. E-CRISPR, CRISPOR and CRISPRscan; Table 1). However, these tools require that the user retrieve the genomic sequences surrounding the variant and select designs that specifically target the variant region after the program sends back all the results. For a bench scientist, this is a time-consuming and error-prone process. For example, when the coding variant is near an exon-intron boundary, the user needs to retrieve the exon sequence as well as the intron sequence and enter these into the program. In addition, the user cannot do batch entries with most of the online tools that take sequence as an input (e.g. E-CRISPR, CRISPOR and CRISPRscan; Table 1). A few command line programs that take sequence as the input were developed for batch design; however, based on our experience, these tools or specific features either do not work or are not easily configured by bench scientists without programing experience (Table 1). In addition, researchers might need features that are missing from current tools, such as an option to target SNPs either together or independently of one another when the SNPs are nearby one another. Moreover, the ability to compare sgRNA designs targeting the same locus in the wild-type and the variant allele in terms of efficiency and specificity would also be very useful when targeting a heterozygous variant.

**Table 1.**
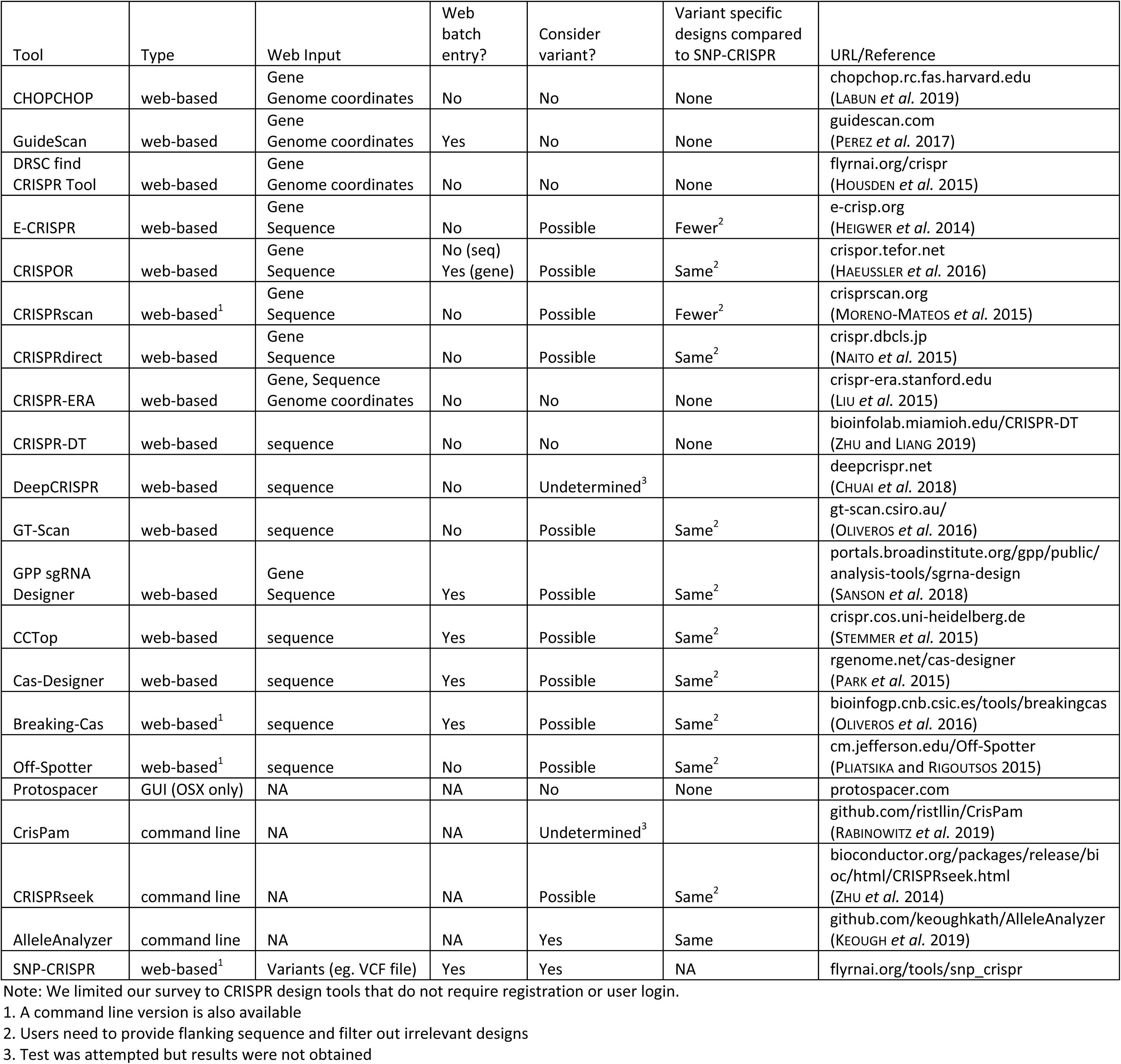
A Survey of CRISPR design tools.

To broaden the application of sgRNA design tools to better accommodate SNPs and small indels, we developed SNP-CRISPR. SNP-CRISPR is a web-based tool that accepts variant annotations as the input and uses rigorous off-target search algorithms to predict the specificity of each target site in the genome for wild-type and variant sequences. SNP-CRISPR offers customized options and allows users to easily and rapidly select optimal variant-specific CRISPR-Cas9 target sequences in genes from a variety of organisms.

## METHODS

### Pipeline development

The SNP-CRISPR pipeline environment is managed using the Conda package and environment management system (ANACONDA 2016). This allows for convenient reproduction of the necessary software dependencies and versions on different machines. The majority of the pipeline logic at SNP-CRISPR is written in Python using Biopython, with some Perl used for the BLAST and efficiency score analysis (COCK *et al.* 2009). Potential off-target loci are evaluated by performing a BLAST search of each design against the species reference genome. An off-target score is assigned based on both the number of hits found in the BLAST results and the number of mismatched nucleotides per off-target hit. Designs are also assigned an efficiency score that was computed using a position matrix; detailed information about the input dataset and algorithm can be found in (HOUSDEN *et al.* 2015). GNU Parallel is used to allow for parallelized computation of designs on different chromosomes and with different parameters for improved performance on multi-core systems (TANGE 2018). The full source code of the pipeline, including instructions for installation and use, is available at https://github.com/jrodiger/snp_crispr.

### Implementation of the web-based tool

The SNP-CRISPR web tool (https://www.flyrnai.org/tools/snp_crispr/) is located at the web site of the Drosophila RNAi Screening Center (DRSC). The back-end is written in PHP using the Symfony framework and the front end HTML pages take advantage of the Twig template engine. The JQuery JavaScript library with the DataTables plugin is used for handling Ajax calls and displaying table views. The Bootstrap framework and some custom CSS is also used on the user interface. Hosting by Harvard Medical School Research Computing makes it possible to provide a web-facing user interface to run the SNP-CRISPR core pipeline on Harvard Medical School’s “O2” high-performance computing cluster. When jobs are submitted from the website, the form parameters and uploaded input file path are passed to a bash script controlling the pipeline, which is then run as a cluster job. When the job is complete, an email is sent to the user with a URL that contains a unique ID used to retrieve the corresponding results.

### Availability

SNP-CRISPR is available for online use without any restrictions at https://www.flyrnai.org/tools/snp_crispr.

The source code for the pipeline, including instructions for installation and use, is available at https://github.com/jrodiger/snp_crispr

## RESULTS

### SNP-CRISPR web tool

The web-based version of SNP-CRISPR provides the functionality of the design pipeline with an easy-to-use interface and interactive results view. Users can select up to 2,000 variants of interest in Variant Call Format (VCF) or a csv file in the provided format, and then upload this file on the SNP-CRISPR homepage. The acceptable variants include single nucleotide changes, small insertions and small deletions. The user then chooses the species and whether to create designs that target each input variant individually or to target all SNPs within each potential sgRNA sequence. When a user submits input, the web logic starts a job on the Harvard Medical School “O2” high-performance computing cluster, using the uploaded file and parameters as input for the pipeline. After the pipeline finishes running, an automated email is sent to the user with a link to a webpage at which the user can view and export results. For a couple of variants, the design pipeline usually takes up to a few minutes and with an input of 2,000 human SNP variants, it takes about half an hour for users to receive the results by email. The result page shows the wild-type and variant designs with corresponding scores in a tabular view that can be sorted by one or more columns. The output table also lists the genome targeting position of each sgRNA and the position of the variant within the sgRNA sequence relevant to the PAM sequence. The variant is shown in lowercase, which can be easily spotted by users. Using the checkboxes in the left-most column, users can opt to export all or only selected rows to an Excel or csv file. Currently, SNP-CRISPR supports reference genomes from human, mouse, rat, fly and zebrafish (Figure 1).

**Figure 1.**
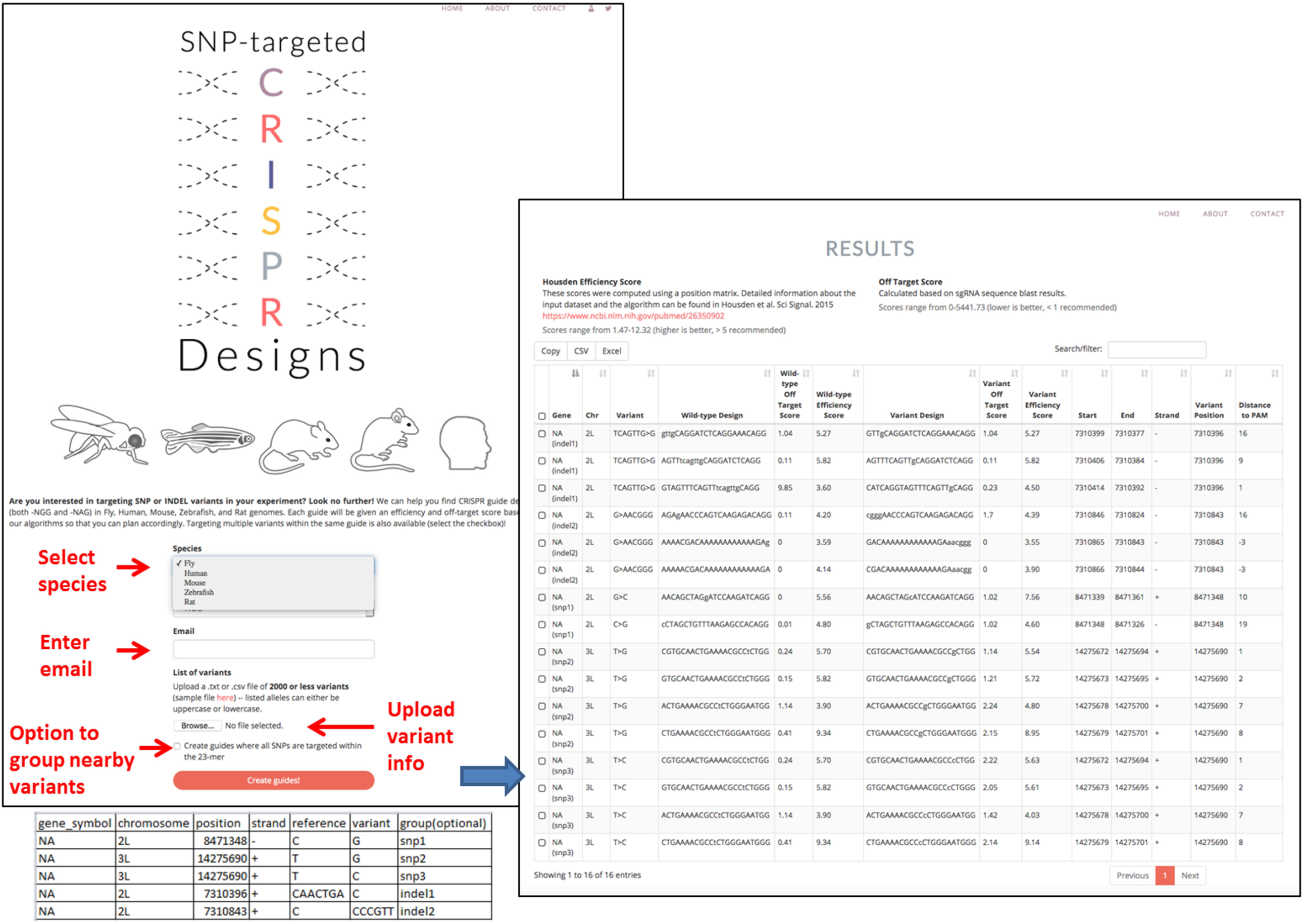
Features of the SNP-CRISPR user interface (UI). Users select the species of interest, enter an email address, upload variant information including the genome coordinates and sequence changes, choose to target nearby variants individually or together, and then submit the job. Usually within half an hour, an email is sent automatically to the user with a link to a results page that displays the designs for wild type as well as mutant alleles, side by side with calculated scores. The mutant base(s) are shown in lower case and the wild type sequence in upper case.

### Computation of potential variant-targeting sgRNAs

Users are required to upload variant information in one of the supported formats including the genome coordinates, the sequence of the reference alleles and the sequence of the variant alleles. First, SNP-CRISPR validates the input reference sequences and will warn users if the submitted reference sequences does not match, which might reflect a different version of the genome assembly being used in the user input vs. SNP-CRISPR. After validation, SNP-CRISPR then re-constructs the template sequence, swapping the reference nucleotide with the variant nucleotide for SNPs, while inserting or deleting the corresponding fragment for indel type variants. Second, SNP-CRISPR computes potential variant-targeting sgRNAs based on availability of PAM sequences in the neighboring region since the presence of a PAM sequence (NGG or NAG) is one of the few requirements for binding. Third, sgRNA designs that contain four or more consecutive thymine residues, which can result in termination of RNA transcription by RNA polymerase III, are filtered out (GAO *et al.* 2018). Cas9 can have off-target activity across the genome and tolerance to mismatches shows significant variance depending on the position within the sgRNA (FU *et al.* 2013; HSU *et al.* 2013). Therefore, for each sgRNA design, SNP-CRISPR computes an efficiency score (HOUSDEN *et al.* 2015) and a specificity score calculated based on BLAST results against the reference genome. All possible sgRNAs are provided to the user along with specificity and efficiency scores, without further filtering; filtering options are available for custom applications based on user needs (Figure 2).

**Figure 2.**
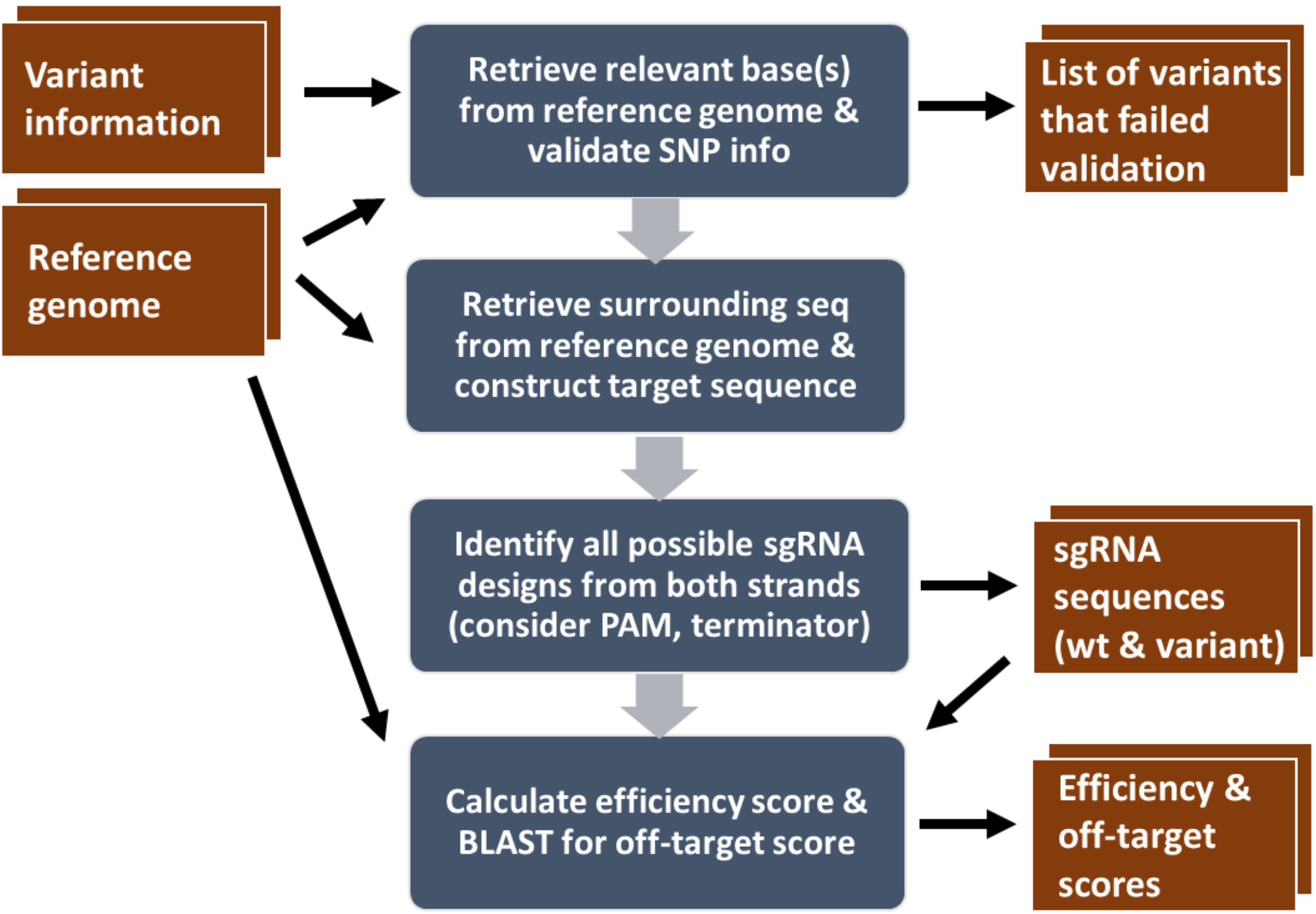
SNP-CRISPR sgRNA design pipeline. Graphic display of the major steps of sgRNA design (blue), and input files and output files for the command line version of the pipeline (red).

To facilitate identification of the best variant-specific sgRNAs, we provide information about both sgRNAs targeting specific variants and sgRNAs targeting the reference sequence in the same region. The efficiency score and an off-target score are provided, and the positions of relevant SNPs or indels in the sgRNA are included so that users can select the most suitable sgRNA or filter out less optimal ones.

The web tool supports up to 2,000 variants per batch while the command line version has no limit with the number of variants and can be used for any annotated genome. The command line version also provides better performance on large inputs when run multi-threaded. For example, a multi-threaded test run was able to process over 1,000 human SNPs per minute on Harvard Medical School’s “O2” high-performance computing cluster. We pre-computed sgRNA designs (NGG-PAM) for all clinically associated SNPs annotated at the Ensembl genome browser (ftp://ftp.ensembl.org/pub/release-97/variation/gvf/homo_sapiens/homo_sapiens_clinically_associated.gvf.gz) using the command line version of the pipeline and the designs can be found at https://github.com/jrodiger/snp_crispr/tree/master/results.

## CONCLUSION

SNP-CRISPR is a unique web tool that designs sgRNAs targeting specific SNPs or indels. SNP-CRISPR is user-friendly and provides all possible CRISPR-Cas9 target sites in a given genomic region with required parameters, allowing users to select an optimal sgRNA. SNP-CRISPR provides not only efficiency scores but also off-target information for sgRNAs targeting sequences with and without SNPs and/or indels of interest in the same genomic region. SNP-CRISPR supports the human reference genome and genomes from major model organisms; namely, mouse, rat, fly and zebrafish. Conveniently, SNP-CRISPR displays the positions of variant nucleotides in each sgRNA region as part of the design output. Moreover, SNP-CRIPSR accepts up to 2,000 inputs per batch for design of large-scale experiments at the website. The command line version has no limit as to the number of variants and can be used for any genome that has been properly annotated. Altogether, SNP-CRISPR improves the ability of researchers to edit SNP or indel-containing loci by facilitating the design of sgRNAs that target specific variants. As such, SNP-CRISPR provides a valuable new resource to the genome editing technology field.

More and more variant data has become available in recent years, and much current research focuses on the biological impact of variants (AMBERGER and HAMOSH 2017; BRAGIN *et al.* 2014; LANDRUM *et al.* 2014; SONG *et al.* 2016), motivating us to develop a variant-centered tool. For instance, a CRISPR/Cas9-based targeting approach has been used to specifically correct heterozygous missense mutations associated with dominantly inherited conditions by including the mutated base in the sgRNA sequence (COURTNEY *et al.* 2016). CRISPR/Cas9-based therapeutic approaches show great promise for permanent correction of genetic disorders in somatic cells. In addition, to facilitate direct research in gene therapy of human diseases, SNP-CRISPR will be valuable for modeling human disease using model organisms. With a vast and growing amount of sequences from different strains of model organisms such as *Drosophila melanogaster*, millions of novel sequence variants have been identified (HUANG *et al.* 2014; WANG *et al.* 2015). However, the biological significance of most of these sequence variants is still unclear. By facilitating design of sgRNAs targeting variant-specific alleles, including at a large scale, SNP-CRISPR makes it more feasible to study these variants systematically.

## ACKNOWLEDGEMENTS

Relevant grant support includes NIH NIGMS R01 GM067761 and P41 GM132087. In addition, C.C. is supported by R21 ES025615, and S.E.M. is supported in part by the Dana Farber/Harvard Cancer Center, which is supported in part by NIH NCI Cancer Center Support Grant P30 CA006516. N.P. is an investigator of Howard Hughes Medical Institute.

